# Genome context influences evolutionary flexibility of nearly identical type III effectors in two phytopathogenic Pseudomonads

**DOI:** 10.1101/2021.11.22.469601

**Authors:** David A. Baltrus, Qian Feng, Brian H Kvitko

## Abstract

Integrative Conjugative Elements (ICEs) are replicons that can insert and excise from chromosomal locations in a site specific manner, can conjugate across strains, and which often carry a variety of genes useful for bacterial growth and survival under specific conditions. Although ICEs have been identified and vetted within certain clades of the agricultural pathogen *Pseudomonas syringae*, the impact of ICE carriage and transfer across the entire *P. syringae* species complex remains underexplored. Here we identify and vet an ICE (PmaICE-DQ) from *P. syringae* pv. maculicola ES4326, a strain commonly used for laboratory virulence experiments, demonstrate that this element can excise and conjugate across strains, and contains loci encoding multiple type III effector proteins. Moreover, genome context suggests that another ICE (PmaICE-AOAB) is highly similar in comparison with and found immediately adjacent to PmaICE-DQ within the chromosome of strain ES4326, and also contains multiple type III effectors. Lastly, we present passage data from *in planta* experiments that suggests that genomic plasticity associated with ICEs may enable strains to more rapidly lose type III effectors that trigger R-gene mediated resistance in comparison to strains where nearly isogenic effectors are not present in ICEs. Taken together, our study sheds light on a set of ICE elements from *P. syringae* pv. maculicola ES4326 and highlights how genomic context may lead to different evolutionary dynamics for shared virulence genes between strains.

## Introduction

Genome fluidity is crucial for survival of bacterial phytopathogens, as selection pressures on the presence and function of specific virulence genes can dramatically change from host to host (Dillon et al., 2019a; de Vries et al., 2020). Much research characterizing gene composition of bacterial pathogens focuses on presence and absence of specific virulence genes, and therefore often extrapolates from lists of loci to predict virulence and evolutionary potential (Baltrus et al., 2011; Dillon et al., 2019b). However, even if virulence genes are conserved across strains of a particular pathogen, differences in genomic flexibility for shared virulence genes could potentiate different evolutionary outcomes between strains under conditions of strong selection. Here we characterize one instance where such differences in genomic context and fluidity for a set of type III effectors affects evolutionary potential for *Pseudomonas syringae*, and we speculate about the ability of such systems to shift evolutionary dynamics for phytopathogens moving forward.

Integrative Conjugative Elements (ICEs) are mobile replicons that blend characteristics of both plasmids and prophage and are well known for their ability to harbor antibiotic resistance genes and other niche-association traits across bacteria (Johnson and Grossman, 2015). Like conjugative plasmids, ICEs contain all genes and pathways required for conjugation between bacterial cells. Like prophage, ICEs contain site specific recombinases that enable recombination into chromosomal locations and repress the genes responsible for conjugation and replication while in this quiescent state. Most importantly, ICEs also contain cargo regions that can house genes and pathways that directly contribute to dramatic changes in bacterial phenotypes (Lovell et al., 2009; Johnson and Grossman, 2015). Not only can these elements modify phenotypes in their current host cells, but the ability of ICEs to transfer throughout populations and across communities through horizontal gene transfer means that cargo genes present on ICEs can rapidly proliferate if they are beneficial. The rise of sequencing technologies enabling closed bacterial genomes has reinforced the importance of mobile elements in genomes and increased awareness of the prevalence of ICEs throughout bacteria.

The phytopathogen *Pseudomonas syringae* (sensu lato) is a recognized agricultural pest for many crops throughout the world, with numerous strains well established as laboratory systems for understanding virulence of plant pathogens *in vitro* and *in planta (Baltrus et al., 2017)*. The presence of a type III secretion system is critical for virulence *in planta* for many strains of this pathogen, and is used to translocate upwards of 40 effectors proteins per strain from the bacterial cytoplasm into plant cells (Dillon et al., 2019a; Laflamme et al., 2020). Once inside the plant cells, effector proteins can disrupt plant immune responses in a variety of ways to promote bacterial growth and infection. The presence of effector proteins can also be monitored by plant immune responses through the action of R-genes, with recognition of effector protein functions leading to an overarching immune reaction termed effector triggered immunity (ETI)) (Collmer et al., 2000; Jones and Dangl, 2006; Jayaraman et al., 2021). Triggering of the ETI response can quickly shut down nascent infections in resistant plant cultivars and has thus formed the basis of future plans to engineer durable crop resistance to infection through genetic modification and selective breeding (Dong and Ronald, 2019; Laflamme et al., 2020). Thus, type III effectors sit at an evolutionary inflection point where they can be highly beneficial for bacterial growth in some host backgrounds and highly detrimental in others.

The presence of virulence genes, including type III effectors, on plasmids and consequent movement across strains can dramatically alter trajectories of virulence evolution for *P. syringae* on different host species (Cazorla et al., 2002; Schierstaedt et al., 2019). Plasmids can be lost or modified if effector proteins are recognized by a potential host, which could enable infections to proceed at a population level despite the ETI response (Grant et al., 2006; Bardaji et al., 2019). Plasmids are not the only element within *P. syringae* genomes displaying plasticity, though. For instance, the effector AvrPphB (aka HopAR1) triggers HR in bean plants that contain the R-gene RPS5 (Qi et al., 2014). AvrPphB is found on an genomic island in certain strains of *P. syringae* pathovar phaseolicola, and under selective pressure from ETI responses bacteria with the island excised from the chromosome and maintained independently as an episome rapidly dominate the population (Godfrey et al., 2011; Neale et al., 2018). Excision of the genomic island containing AvrPphB prevents ETI because this Avr gene is downregulated under episomal replication, but enables this gene to be maintained within a population for infection of host plants where it may be beneficial. ICE elements have also been identified and characterized in *P. syringae* pathovar actinidae strains where their presence/absence is one of the most glaring differences between closely related strains and where they contribute to large scale differences in strain metabolism and resistance to antibacterial compounds (Colombi et al., 2017; McCann et al., 2017; Poulter et al., 2018).

Here we describe how genomic context in a locus encoding an effector protein, HopQ, differentially affects evolutionary flexibility for this genomic island. Recognition of HopQ by the plant R-gene Roq1 triggers ETI in the host plant *Nicotiana benthamiana*, and thus limits growth of strain *P. syringae* pv. tomato DC3000 (*Pto)* in this host (Wei et al., 2007; Schultink et al., 2017). We characterize the genomic context for *hopQ* and other linked virulence genes in both *Pto*DC3000 and another pathogen, *Pseudomonas syringae* pv. maculicola ES4326 (*Pma*ES4326). We demonstrate that this and other effectors are differentially present in an ICE element in *Pma*ES4326 but not in *Pto*DC3000, and confirm that recognition of HopQ also limits growth of *Pma*ES4326 in *N. benthamiana*. These differences are somewhat surprising because at a broad scale these regions appear syntenic. We further show how this differential genomic context allows for differential genomic plasticity for *hopQ* between these two strains, by demonstrating that passage of strains in *Nicotiana benthamiana* leads *hopQ* to be more readily lost in *Pma*ES4326 than in *Pto*DC3000. Thus, we directly demonstrate how genomic context for homologous virulence genes in two closely related pathogens can contribute to different evolutionary trajectories for these strains.

## Methods

### Bacterial strains and culturing conditions

*E. coli* was routinely cultured at 37°C in LB (Lysogeny broth; per 1 L = 10 g tryptone, 5 g yeast extract, 10 g NaCl, pH 7.5 + with 15 g agar for solidified media). *P. syringae* was routinely cultured at 25°C or 28°C in either LM (LB modified; per 1L = 10.0 g tryptone, 6.0 g yeast extract, 2.0 g K_2_HPO_4_·3H_2_O, 0.6 g NaCl, 0.4 g MgSO_4_·7H_2_O, with 18 g agar for solidified media), KB (King’s B; per 1 L = 20.0 g proteose peptone 3, 0.4 g MgSO_4_·7H_2_O, glycerol 10 mL, 2.0 g K_2_HPO_4_·3H_2_O, with 18 g agar for solidified media) or KBC (KB amended with boric acid 1.5 g/L and cephalexin 80 mg/L). Liquid cultures were incubated with shaking at 200 rpm. Where appropriate, media was augmented at final concentration with rifampicin (Rf) 40-60 μg/mL, gentamicin (Gm) 10 μg/mL, kanamycin (Km) 50 μg/mL, spectinomycin (Sp) 50 μg/mL, and diaminopimelic acid (DAP) 200 μg/mL for liquid media, 400 μg/mL for solid media for the growth of DAP-auxotrophic *E. coli* strains.

### DNA manipulation

Plasmid DNA was routinely purified using the GeneJet plasmid miniprep kit (Thermo Fisher). PCR was conducted with Phusion HiFi polymerase (Thermo Fisher). PCR/reaction cleanup and gel extraction were conducted with the Monarch PCR and DNA cleanup kit and Monarch DNA gel extraction kits (NEB). *E. coli* transformation was conducted by either by preparing competent cells using the Mix and Go! *E. coli* transformation kit (Zymo Research) or standard electro-transformation protocols. *Pma*ES4326 strains were transformed with pCPP5372hopQ (Schultink et al., 2017) plasmid DNA via electro-transformation after washing and concentration of overnight liquid cultures with 300 mM sucrose (Choi et al., 2006). Restriction enzymes, T4 ligase, and Gibson Assembly Mastermix were purchased from NEB. Oligonucleotide primers were synthesized by IDT. Commercial molecular biology reagents were used in accordance with their manufacturer’s recommendations.

To create the site-specific Tn*7* 3xmCherry labeling vector pURR25DK-3xmcherry, pURR25 (Teal et al., 2006) was first digested with PstI and recircularized with T4 ligase to remove the *nptII* Km^R^ marker gene creating pURR25DK. To replace the *gfp* gene in pURR25DK, the 3xmCherry cassette was PCR amplified from pGGC026 (pGGC026 was a gift from Jan Lohmann (Addgene plasmid # 48831; http://n2t.net/addgene:48831; RRID:Addgene_48831) using primers bko374 (5’**ACATCTAGAATTAAAGAGGAGAAATTAAGCATGGTGAGCAAGGGCGAGGAGGATAACATG 3’) and** bko375 (5’ **CAGGAGTCCAAGCTCAGCTAATTAAGCTTACTTGTACAACTCATCCATACCACCTGT TGA 3’)** to introduce 30 bp 5’ overlaps corresponding to the *gfp* flanking regions. Gibson assembly was used to join the 3xmCherry PCR amplicon with pURR25DK backbone digested with *Bse*RI and partially digested with *Hind*III.

### Bacterial conjugation and creation of mutant strains

Conjugations were performed by mixing 15 μL of 5-fold concentrated, washed, overnight LM liquid cultures of each parent strain and co-culturing at 28°C overnight on sterile nitrocellulose membranes on either LM or LM+DAP plates (for conjugations with DAP-auxotrophic *E. coli* donor strains). Tn*7* transposition conjugations always included the *E. coli* RHO3 pTNS3 Tn*7* transposase helper strain as a third parent. For all conjugation experiments cultures of each parent strain were included separately as controls. *Pma*ES4326 merodiploid exconjugants of pCPP5729 (pK18msGmΔ*hopQ1)* were recovered on LM Km (Kvitko et al., 2009). Resolved Pma ES4326 pCPP5729 merodiploids were recovered via counter-selection on LM Rf +10% sucrose and sucrose resistant clones were screened for kanamycin sensitivity by patch plating indicating the loss of the pK18ms plasmid backbone (Kvitko and Collmer, 2011). Derivative *att*Tn*7-*3xmCherry transposant strains were recovered on LM Sp and pink clones were selected after 4 days incubation at 4°C and restreaked to isolation. To test the native mobility of PmaICE-DQ, conjugation was conducted as described above with the *Pma* ES4326 pCPP5729 merodiploid as the donor parent and *att*Tn*7-*3xmCherry derivatives of *Pto* DC3000 *Pma* ES4326 ΔPmaICE-DQ strains (*Pma*ES4326ΔDQ3 and *Pma*ES4326-C-LA-P5-20-1) as recipients. PmaICE-DQ exconjugants were recovered on LM Rf Sp Km. Conjugation frequency was calculated as the number of Sp^R^Km^R^ colonies recovered per recipient CFU as determined by dilution plating.

### *Nicotiana benthamiana* growth, inoculation and bacterial passage assays

*Nicotiana benthamiana*, WT LAB accession, and *roq1-1* (Qi et al., 2018) were grown in a Conviron Adaptis growth chamber with 12 h light (125 μmol/m^2^/s) at 26°C and 12 h dark at 23°C. Plants were used at 5-7 weeks post germination. To prepare inoculum, *P. syringae* cultures were recovered from fresh KB plate cultures, resuspended in 0.25 mM MgCl_2_, standardized to OD_600_ 0.2, and serially diluted 10,000X to ∼3×10^4 CFU/mL. Cell suspensions were infiltrated with a blunt syringe into either the 3rd, 4th or 5th leaves. Infiltrated spots were allowed to dry fully and then plants were covered with a humidity dome and kept at 100% humidity for 6-8 days to allow symptoms to develop. At end point, leaves were photographed to document symptoms and four 4 mm leaf punches (∼0.5 cm^2 total) were collected with a biopsy punch from each infiltrated area. Discs were macerated in 0.1 mL 0.25 mM MgCl_2_ using an Analytik Jena SpeedMill Plus homogenizer and the bacterial CFU/cm^2^ leaf tissue was determined by serial dilution spot planting on LM Rf.

For *P. syringae* passaging in *N. benthamiana*, single colonies of *Pto* DC3000 WT and *Pma*ES4326 were inoculated into LM Rf liquid cultures. Samples of the initial cultures (P0) were cryo-preserved at −80C in 15% final strength sterile glycerol. The liquid cultures were diluted 1000X in 0.25 mM MgCl_2_ to approximately 5×10^5^ CFU/mL prior to syringe infiltration into three leaf areas establishing 3 lineages (A, B, C) each for Pto DC3000 and Pma ES4326. Tissue samples were collected 6-7 days post inoculation and processed as described above. Bacteria cultured from the 10^−1^ tissue macerate dilutions of each lineage were directly scraped from the dilution plate with an inoculation loop and suspended in 1 mL 0.25 mM MgCl_2_. These suspensions were then sub-cultured 5 μL into 5 mL LM Rf and diluted as described above to create inoculum for the next passage. This passaging scheme was repeated five times and samples of both the post-passage recovered bacteria (P1) and the corresponding sub-cultured bacteria (P1c) were cryo-preserved for each lineage and each passage. To screen for changes in *N. benthamiana* disease compatibility with passaged strains in a medium-throughput format, bacteria from the P5 cryo-preserved samples were streaked to isolation on KBC plates and isolated “P5” colonies were cultured in 200 μL of LM Rf in sterile 96 well microtiter plates along with Pto DC3000, *Pto* DC3000Δ*hopQ, Pma* ES4326 and *Pma*ES4326ΔDQ3 control strains. Cultures were serially-diluted 5000X in 0.25 mM MgCl_2_ to ∼3×10^4 CFU/mL and inoculated into *N. benthamiana* leaves as described. Inoculum concentration was verified via serial dilution spot plating. Inoculated areas were monitored for disease symptoms compared to their respective control strains over the course to 6-8 days and CFU/cm^2^ leaf tissue was determined for each strain as described above. Isolated colonies of strains that displayed bacterial load and symptoms comparable to their respective Δ*hopQ* backgrounds were subcultured from the dilution spot count plates and retained.

### Genome sequencing and Assembly

For each strain with a genome sequence reported herein, a frozen stock was streaked to single colonies on King’s B (KB) agar plates, at which point a single colony was picked to 2mL KB liquid and grown overnight on a shaker at 27°C. Genomic DNA was extracted from these overnight cultures using a Promega (Madison, WI) Wizard kit and including the optional RNAse step. Each strain was sequenced using multiple technologies, and in each case independent DNA isolations were used to prepare libraries for different sequencing platforms. Illumina sequencing for strains was performed by MiGS (Pittsburgh, PA) using their standard workflow. As described in (Baym et al., 2015), this workflow uses an Illumina tagmentation kit for library generation, followed by sequencing on a NextSeq 550 instrument with 150-bp paired-end reads. Trimmomatic was used for adaptor trimming (Bolger et al., 2014). For nanopore sequencing for strain *Pma*ES4326-D, a library was prepared from unsheared DNA using the Rapid sequencing kit (SQK-RAD004) and sequenced on a Flongle flowcell. For nanopore sequencing for strains *Pma*ES4326-C,*Pma*ES4326ΔDQ3, and *Pma*ES4326-C-LA-P5-20-1, each library was prepared using unsheared DNA as an input to the LSK109 ligation sequencing kit and was sequenced on R9.4 MinION flowcells. Nucleotide bases were called during sequencing using Guppy v3.2.6 in Fast-Mode. All genomes were assembled using Unicycler v.0.4.8 (Wick et al., 2017). The public facing genomes for *Pma*ES4326-D and *Pma*ES4326-C were annotated using PGAP (Tatusova et al., 2016). Genomes for *Pma*ES4326ΔDQ3, and *Pma*ES4326-C-LA-P5-20-1 found in Figshare were annotated using Prokka v 1.14.6 (Seemann, 2014). Default parameters were used for all software unless otherwise specified.

## Results

### Complete Genome Sequences for Multiple Isolates of *Pma*ES4326 Highlights Genome Flexibility for this Strain

We previously reported a draft genome sequence for an isolate of *Pma*ES4326 acquired from the lab of Jeff Dangl (Baltrus et al., 2011), and our first goal was to generate a complete genome sequence for this strain (referred to herein as *Pma*ES4326-D, Table 1). We used MiGS (Pittsburgh, PA) to generate Illumia reads for *Pma*ES4326-D, and their workflow generated a total of 3,284,990 paired reads and 418Mb (∼64x coverage) of sequence. We also independently isolated genomic DNA and sequenced using an Oxford Nanopore MinION to generate 32,291 reads for a total of 465Mb (∼71x coverage) of sequence with a read N50 of 30,656 bp. Assembly of these reads resulted in a complete circular chromosome and four separate plasmids. Notably, our isolate of *Pma*ES4326-D contains three previously reported plasmids (pPma4326A, pPma4326B, pPma4326E) but also appears to have lost two different plasmids (pPma4326C and pPma4326D) first reported as present in this strain by Stavrinides et al., 2004. The assembly for this strain appears to contain an additional ∼350kb plasmid that was not reported by Stavrinides et al. and which we name pPma4326F following earlier naming conventions.

**Table 1:** Sequencing of *Pma*ES4326 Strains

| Table 1: Sequencing of *Pma*ES4326 Strains |  |  |  |  |  |
| --- | --- | --- | --- | --- | --- |
| Strain | Reads (Illumina/Nanopore) | Total bp Sequenced Mb (Illumina/Nanopore) | Read N50 Nanopore bp | Total Sequencing Depth (Illumina/Nanopore) | SRA Accession (Illumina/Nanopore) |
| *Pma*ES4326-D | 3,284,990 / 32,291 | 418.372 / 465.212 | 30,656 | 64x / 71x | SRR15988725 / SRR15988724 |
| *Pma*ES4326-C | 3,757,284 / 55,845 | 485.469 / 700.483 | 23,669 | 74x / 107x | SRR15988571 / SRR15988568 |
| *Pma*ES4326ΔDQ3 | 2,479,212 / 15,275 | 331.276 / 178.118 | 31,491 | 51x / 27x | SRR15988570 / SRR15988567 |
| *Pma*ES4326-C-LA-P5-20-1 | 1,372,720 / 9,934 | 189.971 / 146.897 | 31,020 | 29x / 22x | TBD / SRR15988569 |

Given the absence of two plasmids from *Pma*ES4326-D, we sought to generate a genome assembly from a different lab isolate, acquired from the lab of Alan Collmer referred to here as *Pma*ES4326-C (Table 1). We used MiGS (Pittsburgh, PA) to generate Illumia reads for the Collmer lab version of *Pma*ES4326, and their workflow generated a total of 3,757,284 paired reads and 485Mb (∼74x coverage) of sequence. We also independently isolated genomic DNA and sequenced using an Oxford Nanopore MinION to generate 55,845 reads for a total of 700Mb (∼107x coverage) of sequence with a read N50 of 24,669bp. Hybrid assembly of all read types resulted in a single chromosome that appears nearly complete but was not circularized by Unicycler. However, this assembly does contain all predicted circular plasmids as well as pPmaES4326F.

### An ICE Element Hotspot In *Pma*ES4326

Genomic inspection of the region containing type III effectors *hopQ* and *hopD* in strains *Pma*ES4326-C and *Pma*ES4326-D indicate that this area is a potential hotspot for genomic plasticity. Specifically, in a region bordered by *clpB* and the type III effector *hopR*, gene content characterization strongly suggests the presence of two independent integrative conjugative elements (ICEs, Figure 1). Both ICEs are approximately 70kb in length, and are found adjacent to tRNA loci encoding a proline anticodon. Moreover, roughly 70% of each element is composed of sequences with >95% nucleotide similarity and which encode many of the structural genes predicted to be involved in ICE proliferation and integration. These conserved genes include predicted integrases/recombinases, pilus proteins and ATPases, regulator proteins, a topoisomerase (*topB*) and helicase, chromosome partitioning proteins (*parB*), DNA coupling proteins (*traD*), an NTPase (*traG*), and numerous loci annotated as “integrative conjugative element proteins”.

**Figure 1.**
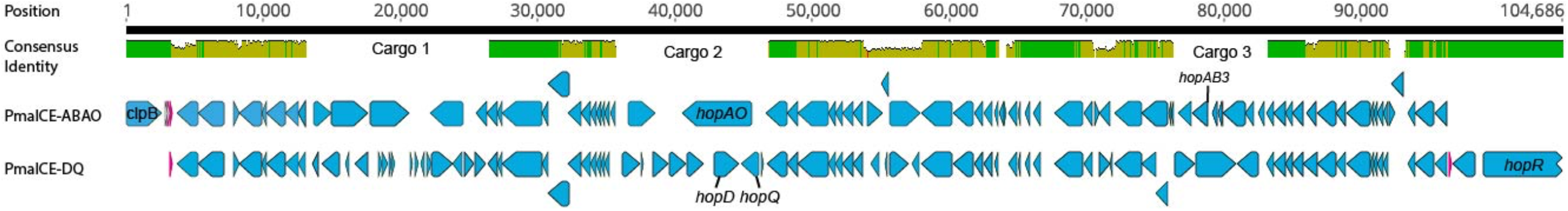
Comparison of PmaICE-ABAO and PmaECE-DQ. We aligned nucleotide sequences from the pair of potential ICEs found within the *clpB-hopR* region in strain *Pma*ES4326 against each other, and visualize the results here. The top line of the figure displays overall nucleotide identity between these two ICEs, with green bars representing conserved nucleotides and yellow representing slight to modest divergence. If no colored bars on this top line, the sequences are completely divergent, and we highlight the three potential cargo regions that differ between these ICE elements. Predicted loci for each ICE are shown on the lines immediately below the nucleotide identity comparison. *Pma*ICE-ABAQ is positioned immediately adjacent to *clpB* in the *Pma*ES4326 genome while *Pma*ICE-DQ is positioned immediately adjacent to *hopR* in the same genome. Each predicted locus is represented in the figure in blue, while predicted tRNA loci are represented in magenta. We labelled all identified type III effector loci within these regions as well as *clpB*.

Despite relatively high sequence similarity across these two closely related ICEs positioned successively in the genome of *Pma*ES4326, gene comparisons suggest the presence of three distinct cargo regions that clearly differentiate these elements (Figure 1). The first ICE contains type III effector proteins *hopAO* and *hopAB3-1* (*hopPmaL*) in two different predicted cargo regions, and so we name this region *Pma*ICE-ABAO. The second ICE contains loci for type III effectors *hopD* and *hopQ* as well as the lytic transglycosylase *hopAJ1*, and so we name this ICE *Pma*ICE-DQ. Cargo region one contains four predicted ORFs in *Pm*aECE-ABAO and fourteen predicted ORFs in PmaICE-DQ. Cargo region two contains two predicted ORFs in PmaICE-ABAO and seven predicted ORFs in PmaICE-DQ, including the effectors *hopAO, hopD*, and *hopQ*. Cargo region three contains 6 predicted ORFs in PmaICE-AO and two in PmaICE-DQ. Aside from the type III effectors, many of loci potentially encoded by these cargo regions are annotated as hypothetical proteins or as parts of IS elements and transposons.

### Differential contexts for the genomic island containing *hopD* and *hopQ* across *Pma*ES4326 and *Pto* DC3000

The type III effectors HopD and HopQ are nearly sequence identical in two phytopathogens commonly used as laboratory models for studying *P. syringae* virulence, *Pto*DC3000 and *Pma*ES4326 (Baltrus et al., 2011). Furthermore, these effectors are found in roughly the same genomic context in the two strains, bordered on one side by *clpB* and on the other by the type III effector *hopR* (Figures 1 and 2). Broad comparisons between *Pto*DC3000 and *Pma*ES4326 further suggest that these effectors form a genomic island along with a third effector (*hopR*) across these two strains (Figure 2), an island which has been referred to as cluster IV in in the *Pto*DC3000 genome (Kvitko et al., 2009). Moreover, the size of the region between *clpB* and *hopR* in *Pto*DC3000 is roughly 70kb, which closely approximates the size of each ICE in *Pma*ES4326. However, while we note above that this region is likely one of high genomic plasticity in *Pma*ES4326, the region containing *hopDQ* in PtoDC3000 lacks many of the hallmarks associated with either PmaICE-DQ or PmaICE-ABAO. While this region for PtoDC3000 does contain subtle hints that suggest the presence of past ICE integration, such as a recombinase syntenic with and similar to those found in the *Pma*ES4326 ICEs, numerous parts of this region in *Pto*DC3000 are quite divergent from that in *Pma*ES4326 (Figure 2). Specifically, numerous ICE-associated proteins appear to be absent from this region in *Pto*DC3000 (Figure 2), and numerous insertion sequence elements and transposase genes have instead proliferated. Such diversity renders accurate evaluation and comparison of evolutionary histories between *Pto*DC3000 and *Pma*ES4326 very challenging at the moment.

**Figure 2.**
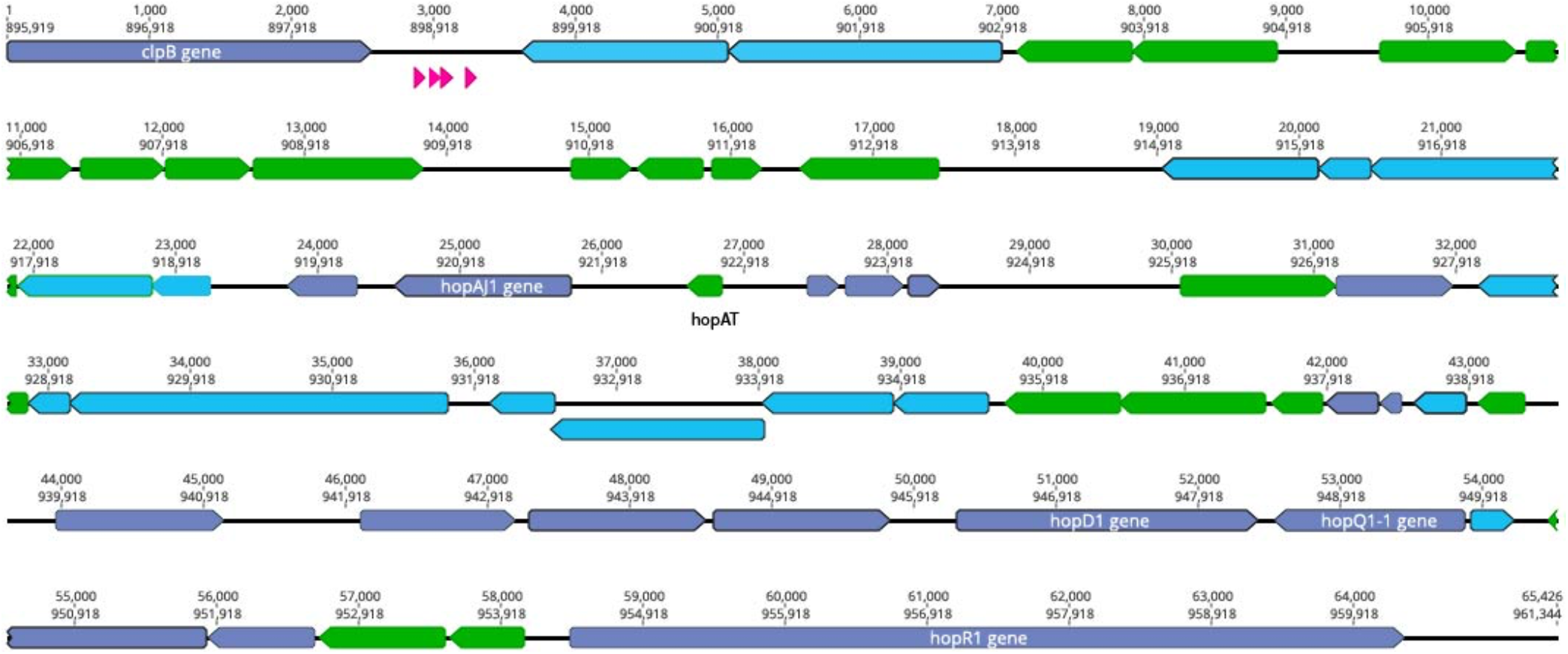
The *clpB-hopR* Region in *Pto*DC3000. Here we visualize the region between *clpB* and *hopR* from the *Pto*DC3000 genome, and compare this region to both predicted ICE elements from *Pma*ES4326. Each predicted locus is colored as purple (present in one of the *Pma*ES4326 ICE elements), blue (present in both *Pma*ES4326 ICE elements as per nucleotide identity), and green (only present in *Pto*DC3000). Predicted tRNA loci are shown in magenta. We also show two nucleotide positions within this region: top number is from the start of *clpB* and the bottom number is from the start of the genome (from *dnaA*). We label all identified type III effector loci as well as *clpB*.

### An evictable ICE Containing HopQ and HopD in *Pma*ES4326

ICEs are categorized by their ability to cleanly excise from the genome, and we confirmed the prediction that the type three effectors *hopQ* and *hopD* are contained in an excisable region in *Pma*ES4326 in two distinct ways. As a first step, we characterized the size of the region excised during intentional creation of a *hopQ-* mutant. To do this, we generated a merodiploid strain in which plasmid pCPP5729 was recombined into a region adjacent to *hopQ*, and then selected for resolution of the merodiploid through plating on sucrose. Presence of *sacB* in plasmid pCPP5729 renders the merodiploid sensitive to killing by sucrose. Notably, this merodiploid also contains regions sequence identical to those surrounding *hopQ* in *Pma*ES4326 and so we originally expected that the *hopQ* gene could be locally deleted through RecA-dependent homologous recombination. We isolated sucrose resistant isolates after plating this merodiploid, and identified strains that were neither WT reverants nor clean *hopQ* deletions by PCR genotyping using previously validated primers. We refer to the focal one of these strains hereafter as *Pma*ICEΔDQ3. Interestingly, while no clean *hopQ* deletion mutants were identified after sucrose counter-selection of merodiploid strains, WT revertants were. We then performed whole genome sequencing to confirm whether the PmaICE-DQ genomic region was lost under these selective conditions in *Pma*ICEΔDQ3. As one can see in Fig. 3, *hopQ* and the surrounding regions that are predicted to be part of PmaICE-DQ have been lost in this strain, confirming that this region can be cleanly and completely excised from the chromosome in a manner consistent with the action of site-specific recombinases contained by ICEs. Searches of both the raw Illumina and Nanopore reads for remnants of *hopQ* or *hopD* yielded no hits despite extensive depth (∼51x for Illumina reads and ∼27x for Nanopore reads), which suggests that PmaICE-DQ was fully lost and not retained as an episome in *Pma*ICEΔDQ3 (data not shown).

**Figure 3.**
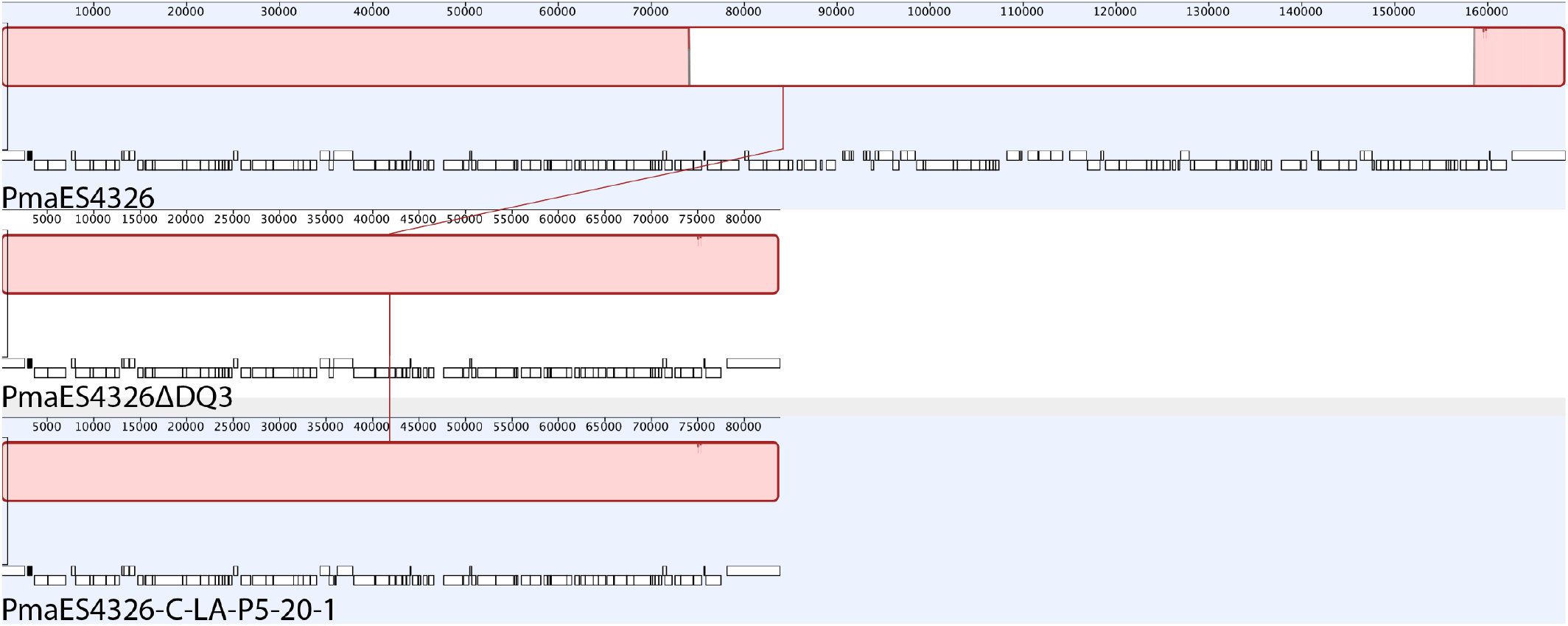
Identical Deletions of PmaICE-DQ following laboratory selection and passage under selective conditions *in planta*. We compared genomic regions bordering *Pma*ICE-DQ in a strain where the region containing *hopQ* was intentionally deleted (*Pma*ES4326ΔDQ3) as well as a strain that arose after 5 passages *in planta* in *N. benthamiana* (*Pma*ES4326-C-LA-P5-20-1). Mauve alignments of these regions demonstrate that identical evictions of *Pma*ICE-DQ occurred in each strain.

### Loss of PmaICE-DQ Enables Virulence of *Pma*ES436 on *N*. *benthamiana*

The presence of hopQ in *Pma* would be expected to elicit the ETI response in *Nicotiana benthamiana* accesions containing the R-gene Roq1, and a lack of compatible infection and disease. Given this information, we tested whether *Pma*ICEΔDQ3 would gain compatibility with *N. benthamiana*. The ΔPmaICE-DQ strain did in fact gain disease compatibility with *N. benthamiana* in a manner similar to the gain-of-compatibility phenotypes previously observed for PtoDC3000 Δ*hopQ1-1* mutants and *Xanthomonas euvesicatoria* 85-10 Δ*xopQ* mutants (Wei et al., 2007; Adlung et al., 2016). Furthermore, *N. benthamiana* incompatibility could be restored in *Pma*ICEΔDQ3 by *hopQ* complementation *in trans* and PmaES4326 incompatibility was not observed in *roq1-1* CRISPR-edited *N. benthamiana* as shown in Fig. 4 (Qi et al., 2018).

**Figure 4.**
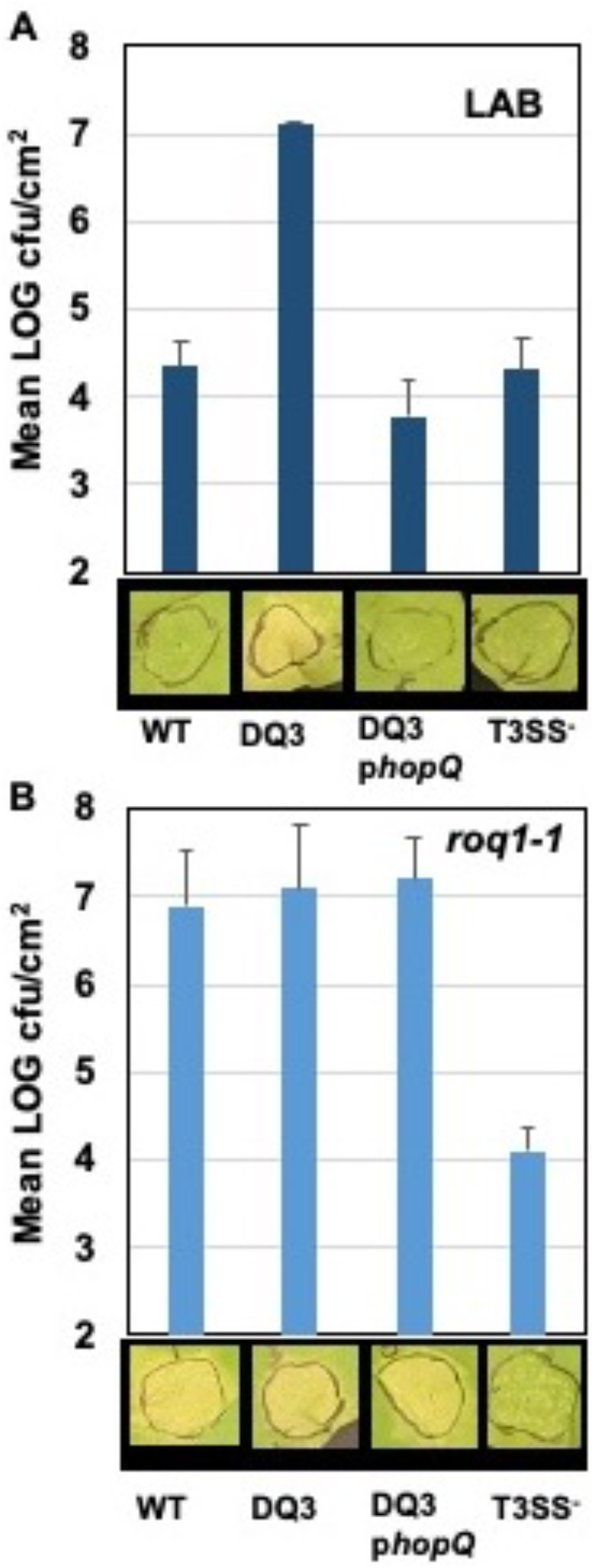
PmaES4326 gains disease compatibility with *N. benthamiana* in the absence of *hopQ*/*Roq1*-mediated ETI. The PmaES4326 WT strain, ΔDQ3 (ΔPmaICE-DQ), ΔDQ3 complemented with pCPP5372hopQ, and the non-pathogenic T3SS-strain *hrcN*::Tn*5* were infiltrated into leaves of the *N. benthamiana* Lab accession (panel A) or the *roq1-1* CRISPR mutant derivative line (panel B) at 3×10^4^ CFU/mL. Eight days post inoculation leaves were photographed to document symptoms and bacterial load from four 4 mm leaf discs from each infiltrated area by dilution plating was determined as CFUs/cm^2^ leaf tissue. Values are the means and standard deviations of LOG transformed CFUs/cm^2^ from three inoculated plants.

### Loss of PmaICE-DQ Under Selection *In Planta*

Observing that we could readily recover strains containing spontaneous evictions of the PmaICE-DQ under the selective pressure of sucrose counter-selection, and that *Pma*ICEΔDQ3 gained disease compatibility with *N. benthamiana*, we tested whether the selective pressure of ETI would also result in recovery of strains with PmaICE-DQ evicted. We inoculated *N. benthamiana* with *Pma*ES4236-C and PtoDC3000 at approximately 5×10^5 CFU/mL establishing three lineages for each strain (Lineage A, LA; Lineage B, LB; Lineage C, LC). Six to seven days post inoculation bacteria from each lineage were recovered and used to create new inoculum to passage the bacteria through *N. benthamiana* a total of five times. Twenty-two isolated clones of each passage 5 (P5) lineage were screened for altered *N. benthamiana* disease compatibility phenotypes. For *Pto*DC3000, none of the 66 P5 clones tested produced disease symptoms distinct from the WT *Pto*DC0000 control. However, for *Pma*ES4326, 7 of 66 (2 LA, 3 LB, 2 LC) P5 clones were able to cause disease symptoms similar to *Pma*ICEΔDQ3 (See Fig. 5). Genome sequencing of *Pma*ES4326-C clone LA-P5-20-1 confirmed the loss of PmaICE-DQ in this strain identical to that observed in the *Pma*ICEΔDQ3 strain recovered after sucrose counter-selection (Table 2, Figure 3).

**Table 2:** Assembly of *Pma*ES4326 Strains

| Table 2: Assembly of *Pma*ES4326 Strains |  |  |  |  |
| --- | --- | --- | --- | --- |
| **Strain** | **Complete Genome** | **Number of Contigs in Assembly** | **Replicons Present** | **Assembly Accession** |
| *Pma*ES4326-D | Yes | 5 | Chromosome (circular), pPmaES4326ABEF | GCA_000145845.2 |
| *Pma*ES4326-C | No | 7 | Chromosome (not circular), pPmaES4326ABCDEF | GCA_020309905.1 |
| *Pma*ES4326ΔDQ3 | Yes | 6 | Chromosome (circular), pPmaES4326ABCF | 10.6084/m9.figshare.17064080 |
| *Pma*ES4326-C-LA-P5-20-1 | No | 7 | Chromosome (not circular), pPmaES4326ABCDEF | 10.6084/m9.figshare.17064080 |

**Figure 5.**
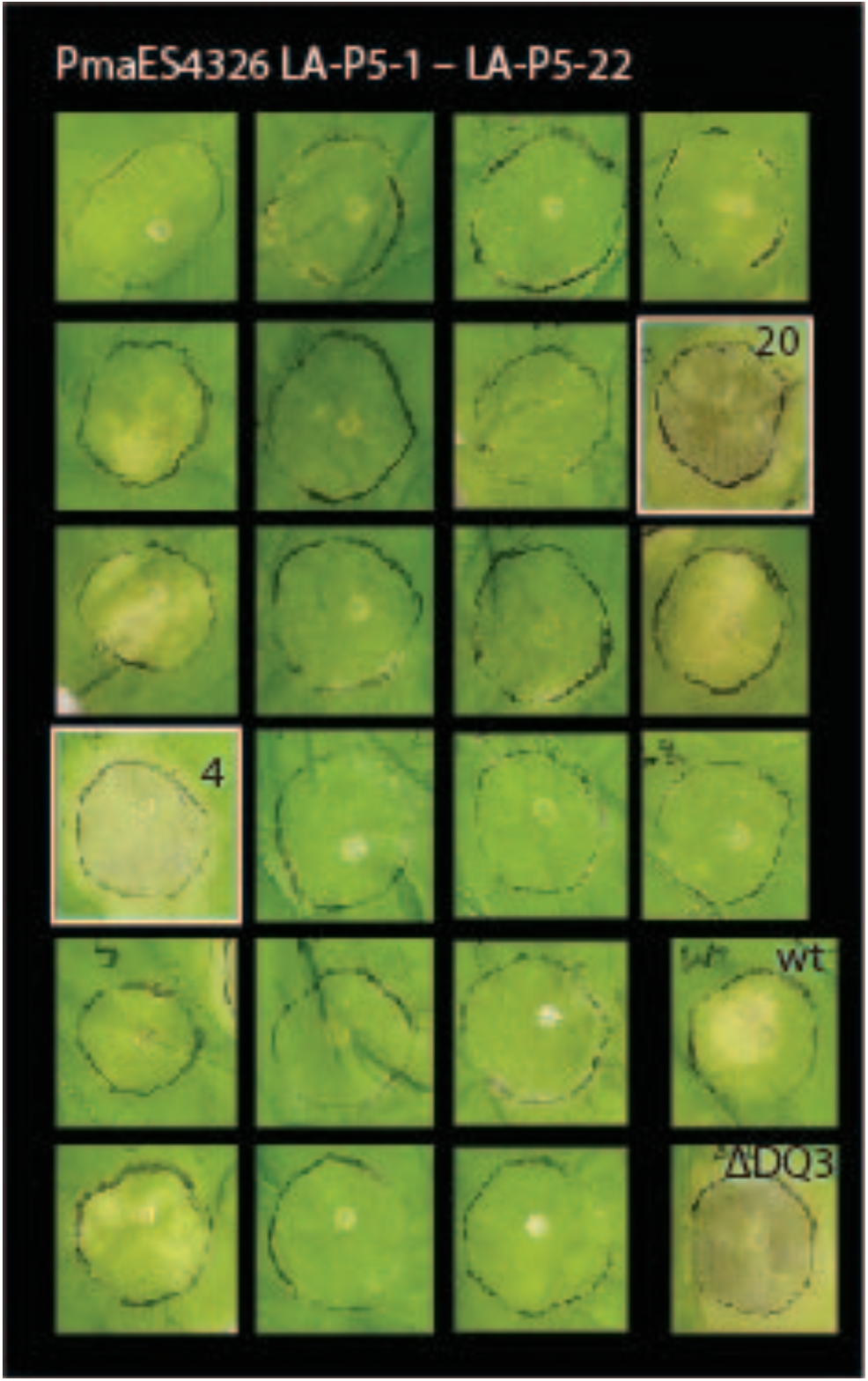
A subset of PmaES4326 isolates passaged through *N. benthamiana* gain disease compatibility. Two isolates out of twenty-two *Pma*ES4326 lineage A (LA) passage 5 (P5) isolates produce disease symptoms in the *N. benthamiana* LAB accession comparable to the ΔDQ3 (ΔPmaICE-DQ) strain. Leaves were photographed eight days post inoculation with 3×10^4^ CFU/mL bacterial suspensions.

### Transfer of PmaICE-DQ Between Strains

Aside from being able to excise from the genome, ICE elements are also categorized by their ability to transfer across bacterial strains, and we therefore tested whether PmaICE-DQ could undergo conjugation to a naive strain. To do this, the PmaICE-DQ merodiploid described above (the precursor to *Pma*ICEΔDQ3) was co-cultured with either *Pto*DC3000 or two independent PmaΔICE-DQ strains (*Pma*ICEΔDQ3 and *Pma*ES4326-C-LA-P5-20-1). After their initial creation, these recipient strains were labeled with a Tn*7* 3xmCherry SpR resistant transposon to differentiate them from the donor strain. Conjugations between these strains were then plated on kanamycin and spectinomycin selective media (Fig. 6A). No kanamycin and spectinomycin resistant mCherry exconjugants were recovered from the *Pto*DC3000 conjugation. However, we were able to recover kanamycin and spectinomycin resistant mCherry+ exconjugants in *Pma*ES4326ΔICE-DQ strains at rates of 2.50×10^−7^ stdev ± 1.23^-7^ and 3.05×10^−7^ stdev ± 1.51^-7^ exconjugants per recipient respectively in derivatives of *Pma*ICEΔDQ3and *Pma*ES4326-C-LA-P5-20-1 (Fig. 6B) which strongly suggests that the PmaICE-DQ element is readily transmissible into *Pma*ES4326 strains lacking PmaICE-DQ.

**Figure 6.**
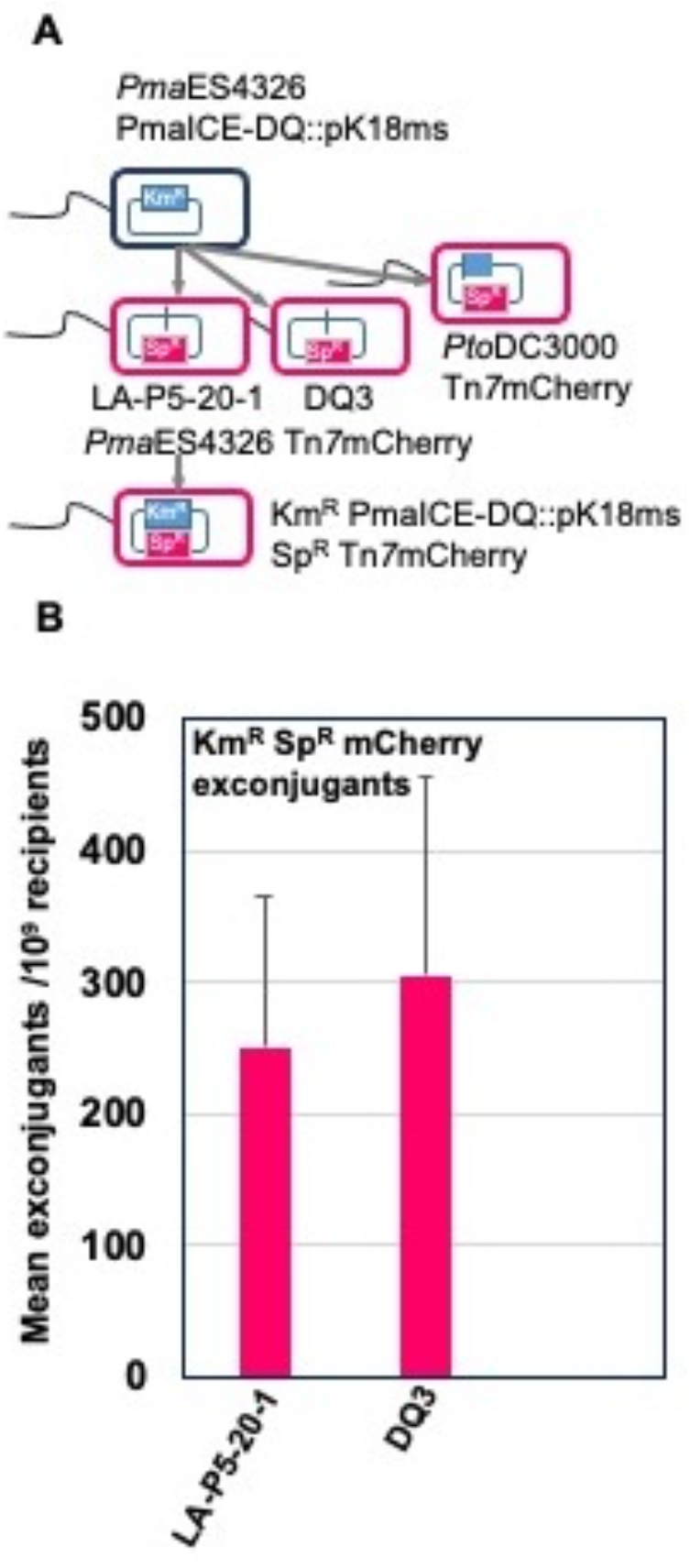
PmaICE-DQ can be readily transferred into PmaES4326 strains that lack the ICE element. (A) Schematic representation of the conjugation strategy for monitoring PmaICE-DQ transfer between strains. In the donor strain, PmaICE-DQ was marked with kanamycin resistance by single crossover integration the pK18ms plasmid. The recipient strains PmaES4326ΔDQ3, PmaES4326-C-LA-P5-20-1 and Pto DC3000 were marked by introduction of a Spectinomycin resistance Tn*7*3xmCherry marker at their *att*Tn*7* sites. Donors and recipients were co-cultured on LM media and PmaICE-DQ mCherry expressing exconjugants were recovered on kanamycin and spectinomycin. (B) PmaICE-DQ exconjugant recovery frequencies/10^9^ recipients were determined based on kanamycin and spectinomycin exconjugants per spectinomycin resistant recipients as determined by dilution plating. Values are means and standard deviations of three biological replicates.

## Discussion

Integrative Conjugative Elements (ICEs) are important drivers of evolutionary dynamics within and across bacterial populations and communities because of their potential to disseminate genes and pathways through horizontal transfer (Johnson and Grossman, 2015). Genes encoded by ICEs often impart phenotypes critical for survival under specific environmental conditions, including antibiotic resistance loci as well as phage defense systems (Botelho and Schulenburg, 2021; LeGault et al., 2021). In the phytopathogen *Pseudomonas syringae*, plasticity of ICE elements encoding genes involved in heavy metal resistance, type III effectors, and carbon metabolism has been identified as the main difference between epidemic strains causing disease across kiwi orchards in New Zealand and with previous outbreaks (Colombi et al., 2017; McCann et al., 2017; Poulter et al., 2018). Despite the accumulation of numerous complete genomes sequences for *P. syringae* and related species and the likelihood of ICEs to impart traits that affect growth in agricultural settings, to date there have been relatively few ICEs identified and vetted for this pathogen.

Placement of genes and pathways in ICEs may be especially important under conditions of fluctuating selection pressures where selection pressures on cargo genes in an ICE switch from positive to negative depending on the environment. One such scenario involves the presence of type three effector genes, which are bacterial proteins delivered from symbionts into eukaryotic host cells and which are critical for pathogenesis of *P. syringae* strains *in planta (Grant et al., 2006)*. Type three effectors are particularly well studied in phytopathogens such as *Pseudomonas syringae*, where they can be of exceptional benefit on some host plants by enabling strains to overcome host immune responses but may also directly trigger immune responses based on the presence of cognate immune receptors on other host genotypes (Dillon et al., 2019a; Laflamme et al., 2020). The presence of type three effectors on ICEs would enable the rapid transmission of these critical virulence genes across strains, while also enabling genes to be rapidly lost from lineages if host genotypes shift such that the effectors are recognized by R gene pathways. Extending this thought, effectors in ICEs that are recognized by hosts are potentially lost more readily through ICE element excision than through mutation (given dedicated excision machinery of the ICEs), which may facilitate adaptation if a strain encounters a resistant host background. While not an extensive experiment, our passage experiment where a subset of *Pma*ES4326 but not *Pto*DC3000 clones becomes compatible with *N. benthamiana* does demonstrate that parameter space exists for such a scenario. Moreover, effectors that are inactivated through mutation can only be reactivated by reversion mutations or through horizontal transfer and acquisition from different strains. In contrast, if effectors are found on ICEs and lost through excision, it’s possible that a small resident pool of strains containing these ICES will remain on other host plants or throughout the environment, facilitating rapid reacquisition of effectors that could be beneficial under different contexts. Lastly, there are a plethora of type III effectors in *P. syringae* that carry out a variety of functions across different host plants and resistance backgrounds (Dillon et al., 2019a). Some of these effectors can be considered “core” and found in syntenic locations across strains while others are more variably present. It’s possible that presence of effectors in ICEs could itself reflect something about the characteristics of these effectors, in that the proteins found as cargo on ICEs may be subject to higher levels of fluctuating selection across hosts than those found in the core set.

HopQ and HopD are often found together on genomic islands in *P. syringae* genomes, and have been associated with adaptation to agriculturally important crop plants (Wei et al., 2007; Baltrus et al., 2011; Monteil et al., 2016). Presence of *hopQ* in an ICE in strain *Pma*ES4326 sets up an interesting scenario because a nearly identical version of this effector is present in a somewhat syntenic position in a relatively closely related strain *Pto*DC3000 except that this effector is not part of an intact ICE. Therefore, this genomic context suggests that *hopQ* would experience more evolutionary flexibility in *Pma*ES4326 than in *Pto*DC3000 due to its presence in an ICE. To test for differences in evolutionary flexibility, we passaged replicate populations of both *Pma*ES4326 and *Pto*DC3000 in a cultivar of *Nicotiana benthamiana* which can recognize and mount an immune response to HopQ. As such, wild type versions of *Pma*ES4326 and *Pto*DC3000 are recognized by this cultivar and trigger an immune response which prevents disease. However, we found that passage of *Pma*ES4326 (but not *Pto*DC3000) from plant to plant rapidly selected for cells of *Pma*ES4326 that didn’t trigger the immune response. Therefore, presence of this type III effector inside of an ICE enables rapid loss of the recognized effector under conditions of negative selection and there is much less flexibility in this loss in a strain where the effector is not present in an ICE.

One other curious feature of the genome of *Pma*ES4326 is that it encodes two distinct but highly similar ICE elements (Figure 1). The configuration is particularly interesting as two distinct ICE elements have integrated successively into the *Pma*ES4326 genome and many of the genes involved in ICE functions are quite conserved between the two. It also appears as though integration of the first element recreated a tRNA with a proline anticodon and that the second ICE element can use this as a further integration point. Indeed, excision of PmaICE-DQ in both strains reported here also deletes one of the predicted proline tRNAs. It may therefore be no coincidence that two ICE elements that are highly similar (but which contain different cargo regions) can integrate successively into the genome as it’s straightforward to imagine that highly specific recombinases would have similar target sequences.

Although not the main focus of this manuscript, we also report on plasticity of the *Pma*ES4326 genome writ large across lab derived strains (Table 2). As the number of complete *P. syringae* genome sequences accumulates, it is becoming more apparent that strain *Pma*ES4326 is notable compared to the rest of the species complex because it contains so many secondary replicons as well as additional elements that contribute to genomic plasticity (Stavrinides and Guttman, 2004; Stavrinides et al., 2012). Notably, many of these plasmids appear to contain genes involved in virulence and so acquisition of these replicons through horizontal gene transfer has likely contributed to the pathogenic ability of this strain across multiple hosts. However, there is a clear downside to the large plasmid repertoire of *Pma*ES4326 as demonstrated by genome sequences reported here. Although we can’t definitively catalogue passage histories, genome sequences from isolates from the Dangl and Collmer labs are likely not diverged by >10 passages total. In this time, the Dangl isolate (*Pma*ES4326-D) has lost two different plasmids that were originally reported by Stavrinides et al. and which are contained in the Collmer lab isolate (*Pma*ES4326-C). Indeed, genome assemblies for the PmaICE-DQ excision strain suggest that this strain has also lost at least one plasmid in addition to the PmaICE-DQ after only 3 additional passages in culture. Lastly, even though this strain was extensively surveyed for plasmids through gel electrophoresis and Sanger sequencing, we report the existence of an additional large plasmid present within all sequenced isolates of this strain.

## Data Availability

Sequencing reads for each of the genomes have been posted in the SRA at accessions listed in Table 1. Genome assemblies for each genome have been posted at either Genbank or Figshare and are listed in Table 2.

## Acknowledgments

We would like to thank Karl Schreiber and Darrell Desveaux for providing *Pma*ES4326 *hrcN*::Tn*5* as well as Alex Schultink and Brian Staskawicz for providing the *roq1-1 N. benthamiana* line. This work is supported in part by the University of Georgia Office of Research as well as the College of Agricultural and Environmental Sciences’ Research Office.

## Notes

https://figshare.com/articles/dataset/Genome_context_influences_evolutionary_flexibility_of_nearly_identical_type_III_effectors_in_two_phytopathogenic_Pseudomonads/17064080

### Competing Interest Statement

The authors have declared no competing interest.

